# Ox-LDL induces a non-inflammatory response enriched for coronary artery disease risk in human endothelial cells

**DOI:** 10.1101/2024.11.19.624299

**Authors:** Jiahao Jiang, Thomas K. Hiron, Anil Chalisey, Yashaswat Malhotra, Thomas Agbaedeng, Chris A. O’Callaghan

## Abstract

Oxidised low-density lipoprotein cholesterol (ox-LDL) is critical in the initiation and progression of atherosclerosis. While excessive atherogenic lipids in the arterial intima can trigger endothelial dysfunction in advanced lesions, the response of endothelial cells to ox-LDL in the early stages of atherogenesis remains unclear. Here, we conducted a comprehensive, genome-wide multi-omics characterisation of the cellular response to ox-LDL in primary human aortic endothelial cells (HAECs). Our results reveal that the exposure of HAECs to ox-LDL leads to pathogenic changes in metabolism, transcriptome and epigenome, but in the absence of a typical inflammatory endothelial phenotype. An integrative analysis implicates the role of AP-1, NFE-2 and CEBP transcription factors in regulating ox-LDL-induced transcription. We further demonstrate that ox-LDL activates endothelial cell migration through the epigenomic rewiring of transcription factor binding. Notably, these ox-LDL-induced dynamic binding sites are enriched for the genetic risk of coronary artery disease, enabling the discovery of the gene-environment interaction of rs62172376 and ox-LDL at the *CALCRL*/*TFPI* locus. Collectively, our findings provide an unbiased understanding of the transcriptional regulation in endothelial cells in response to ox-LDL, together with its interaction with the genetic element of coronary artery disease.

## Introduction

Atherosclerosis is a chronic, multicellular disease characterised by the formation of lipid-rich plaques within the arterial wall, and is the primary underlying cause of coronary artery disease (CAD). Endothelial cells (ECs) constitute the first barrier of the arterial vasculature, and are central to the initiation and the progression of atherosclerosis. In health, ECs tightly control vascular tone, prevent blood clotting and maintain redox balance ^1^; in disease or inflammatory conditions, damage to ECs triggers the recruitment of immune cells to the sub-endothelial space, further exacerbating the inflammatory and necrotic microenvironment within the arterial wall ^2–5^.

Cholesterol-carrying low-density lipoprotein (LDL) particles are retained and chemically modified (e.g., notably by oxidation) in the vessel wall at lesion-prone sites during the initiation of atherosclerosis ^6,7^. Due to their direct exposure to circulating blood, ECs have long been believed to be the first cell type to experience the pathogenic effects of LDL and its derivatives. Specifically, exposure to oxidised LDL (ox-LDL) has been linked to pro-inflammatory endothelial phenotypes, characterised by increased expression of cell adhesion molecules such as ICAM-1 and VCAM-1 ^8–12^. However, contrasting results showing no or context-dependent endothelial activation after ox-LDL exposure were also observed using the same models ^13–17^. Most of these early studies on ox-LDL and ECs were conducted using low-throughput approaches, with a narrow focus on specific pathways or functions hypothesised to be relevant. Therefore, these results have often been inconclusive. We reasoned that a comprehensive, unbiased assessment of the endothelial response to ox-LDL in primary human cells is crucial for a more complete understanding of this pathophysiological process underlying atherosclerosis.

The development of high-throughput multi-omics sequencing approaches enables unbiased, genome-wide investigation of the endothelial response to atherogenic lipids. One of the bioactive components of ox-LDL, oxidised 1-palmitoyl-2-arachidonoyl-sn-glycero-3-phosphocholine (ox-PAPC) has been extensively characterised ^18–20^. In primary human aortic endothelial cells (HAECs), ox-PAPC has been shown to alter the transcriptome ^21^, epigenome ^22^, and metabolome ^23^. While serving as a convenient synthetic alternative to the plasma-derived ox-LDL, ox-PAPC represents only a fraction of the lipid peroxidation products found in ox-LDL, and its pathogenic similarity with ox-LDL has not been systematically evaluated. Given that oxidised lipids do not act in isolation during atherosclerosis, measuring the overall effect of ox-LDL instead of its bioactive components might better reflect its multifaceted atherogenic properties *in vivo*.

Decades of large-scale, genome-wide association studies have revealed a large heritable component of CAD, with more than 330 risk loci reaching genome-wide significance ^24–26^. ECs are significantly associated with the genetic risk of CAD ^27,28^, and a few EC-acting risk variants have been prioritised through fine-mapping studies ^29,30^. Nevertheless, it remains poorly understood which other variants regulate which genes in endothelial cells to modulate disease risk. In particular, it is unclear whether any of these risk genes are regulated in response to atherogenic lipids in endothelial cells, or if they encode proteins that could potentially interact with these lipids.

In the current study, we employ a genome-wide, multi-omics approach to characterise the response of primary human endothelial cells to ox-LDL. We find only limited overlaps in the metabolic and transcriptomic effect between ox-LDL and ox-PAPC; ox-LDL-induced gene expression better aligns with the *in vivo* gene expression signatures in ECs from human atherosclerotic lesions. Using a combination of computational and experimental approaches, we find that ox-LDL triggers endothelial cell migration through epigenomic rewiring of transcription factor binding. In addition, we show that the endothelial response to ox-LDL is associated with the genetic risk of CAD, and prioritise CAD risk variants that overlap dynamic transcription factor binding sites.

## Results

### Ox-LDL accumulation triggers metabolic shift in HAECs

It is well-recognized that endothelial cells play an important role in the transcytosis of lipids from the circulation to the sub-arterial space ^31^. To test whether exposure to ox-LDL results in intracellular accumulation of lipids, primary human aortic endothelial cells (HAECs) were incubated with DiI-labelled ox-LDL and profiled using immunofluorescence microscopy. After 48 hours of DiI-ox-LDL treatment, lipid droplets were clearly visible in the cytoplasm (Figure 1A and 1B), although the number of these droplets is considerably lower compared to what we have previously reported in primary human macrophages ^32^. The dose-dependence of intra-cellular lipid accumulation was further confirmed by flow cytometry (Figure 1C).

**Figure 1.**
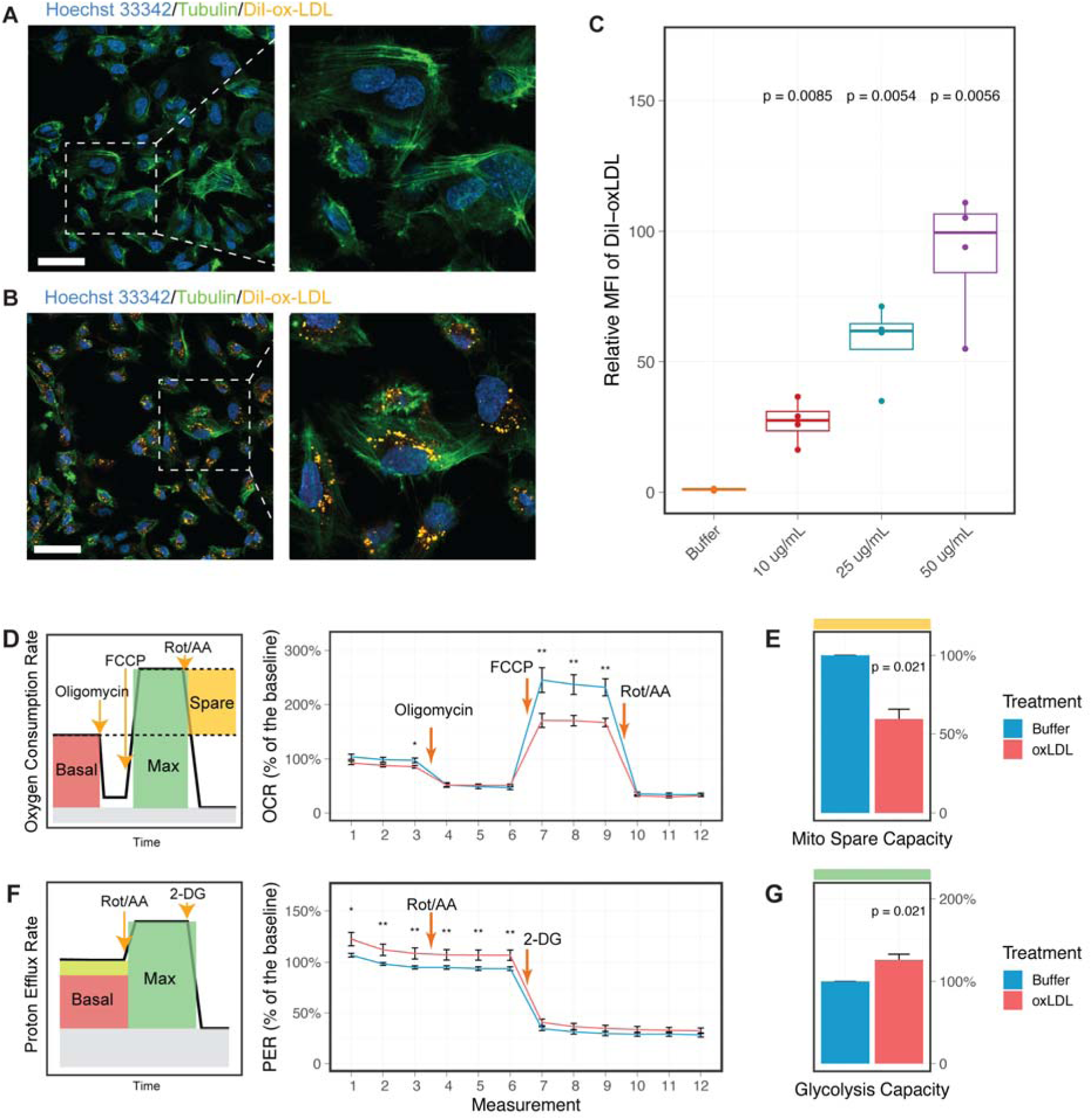
Exposure to ox-LDL induces metabolic stress in primary human aortic endothelial cells (HAECs). **A** and **B**, Representative images from immunofluorescence staining of unexposed (**A**) and ox-LDL-exposed (**B**) HAECs. Cells were treated with 50ug/mL of DiI-labelled ox-LDL or control buffer for 48 hours before imaging. Scale bar: 50um. **C**, Quantification of intracellular DiI signals using flow cytometry in HAECs treated with DiI-ox-LDL for 24 hours at indicated concentrations (n=4 biological replicates). Signals were normalised to the buffer group and shown as the relative mean fluorescence intensity. **D,** Oxygen consumption rate (OCR) of unexposed (blue) and ox-LDL-exposed (red) HAECs, shown as the mean value with the standard error for n=4 biological replicates. Overview of the mitochondrial stress test assay is plotted to the left. **E**, The spare respiration capacity as marked in yellow in **D**, normalised to the buffer group. **F**, Proton efflux rate (OCR) of unexposed (blue) and ox-LDL-exposed (red) HAECs, shown as the mean value with the standard error for n=4 biological replicates. Overview of the glycolytic rate assay is plotted to the left. **G**, The glycolytic capacity as marked in green in **F**, normalised to the buffer group. Seahorse assays were conducted with at least three technical repeats per biological replicate. P-values were calculated using Wilcoxon tests. * p-value< 0.05, ** p-value <0.01.

A previous study has reported that oxidised 1-palmitoyl-2-arachidonoyl-sn-glycero-3-phosphocholine (ox-PAPC), a synthesised bioactive component of ox-LDL, disrupts amino acid metabolism in endothelial cells ^23^. To explore any potential metabolic consequences of ox-LDL exposure, we next measured the cellular respiratory rates and glycolytic rates using Seahorse assays (Figure 1D to 1G).

While the basal respiration rate remains constant, ox-LDL treatment drastically reduced the spare respiration capacity the cells can utilize when the energy demand is high (Figure 1D and 1E). In addition, cells exposed to ox-LDL displayed a significant increase in basal glycolytic rate (Figure 1F and 1G); the addition of rotenone and antimycin A, which inhibits electron transport complexes and thereby shuts down mitochondrial respiration, did not further increase the glycolytic rate (Figure 1F). These results contrast with the metabolic activation effect of ox-PAPC reported previously ^23^, suggesting that ox-LDL induces metabolic stress and pushes the energy balance toward glycolysis.

### Ox-LDL alone does not induce pro-inflammatory endothelial phenotypes

Previous studies have shown that ox-LDL induced endothelium dysfunction, an important pro-atherogenic process that is typically characterized by elevated expression of cellular adhesion molecules such as ICAM-1 and VCAM-1 ^9–12^. However, despite its intracellular accumulation and the metabolic consequences, ox-LDL-exposure was not found to affect the transcript nor protein level of ICAM-1 and VCAM-1 in HAECs (Figure S1A-D).

We noticed that in one of the previous reports ^17^, activation of inflammation was only observed when the lectin-type oxidized LDL receptor 1 (OLR1) was artificially overexpressed in HAECs. Despite being first identified as an endothelial receptor in bovine cells ^33^, *OLR1* is barely expressed in human endothelial cells from either disease-free vasculatures or atherosclerotic plaques (Figure S2). Together, our results clearly demonstrate that OLR1-mediated endothelial activation is unlikely to be a direct consequence of ox-LDL exposure both *in vitro* and *in vivo*.

### Transcriptomic analysis reveals disrupted fatty-acid metabolism upon ox-LDL exposure

To systematically characterize the impact of ox-LDL on gene expression in HAECs, we next performed RNA-seq analysis. After donor and batch effect correction, 304 genes were identified as differentially expressed under the FDR threshold of 0.05, with 144 genes being upregulated by ox-LDL and 160 genes downregulated (Figure 2A). The most upregulated and most downregulated genes are listed in Table 1 and Table 2. Among these genes were several with previously known roles in cholesterol efflux (e.g. *ABCA1*, *ABCC3*) ^34^ and oxidative stress response (e.g. *NQO1*, *HMOX1*) ^35^.

**Figure 2.**
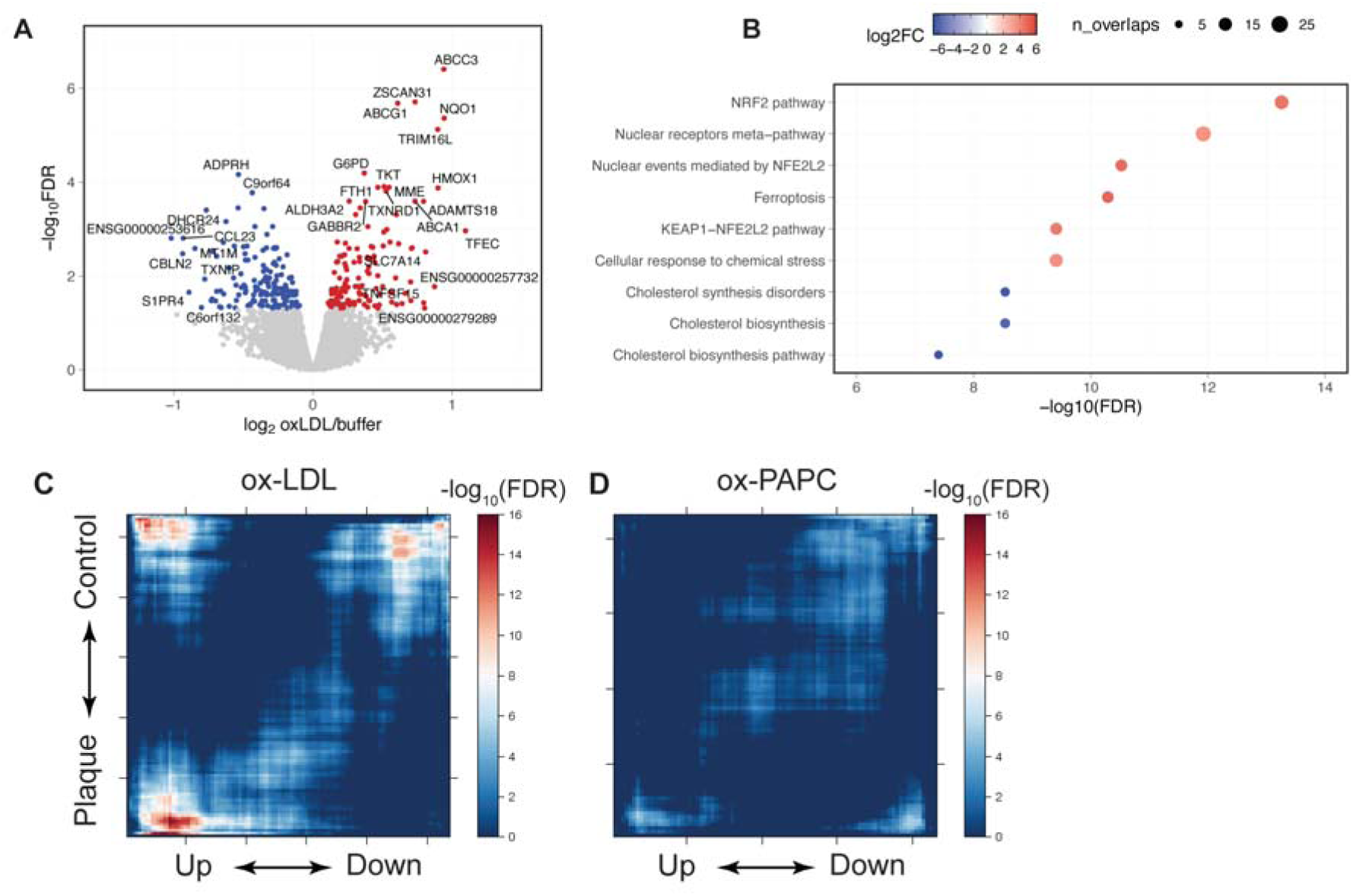
HAECs exposed to ox-LDL display transcriptomic signatures associated with endothelial cells in human atherosclerosis. **A**, Volcano plot showing differentially expressed genes up-regulated (red) and down-regulated (blue) by ox-LDL in HAECs. Genes with FDR>0.05 are plotted in grey. **B**, MsigDB pathways enriched in the differentially expressed genes, ranked by statistical significance. Colour represents log2 fold change of pathway enrichment and size represents the number of overlapped genes in each pathway. **C** and **D**, Rank-rank hypergeometric overlap tests showing similarity between transcriptomic signatures related to plaque ECs (y axis), ox-LDL-treated HAECs (x axis in **C**) and ox-PAPC-treated HAECs (x axis in **D**). Note that genes were ranked by statistical significance of differential expression tests, so only the ends of x or y axis represent significant differentially-expressed genes. The colour denotes the significance of each hypergeometric test (-log_10_FDR), measuring the extent of overlap between the two gene set on a sliding basis.

**Table 1.**
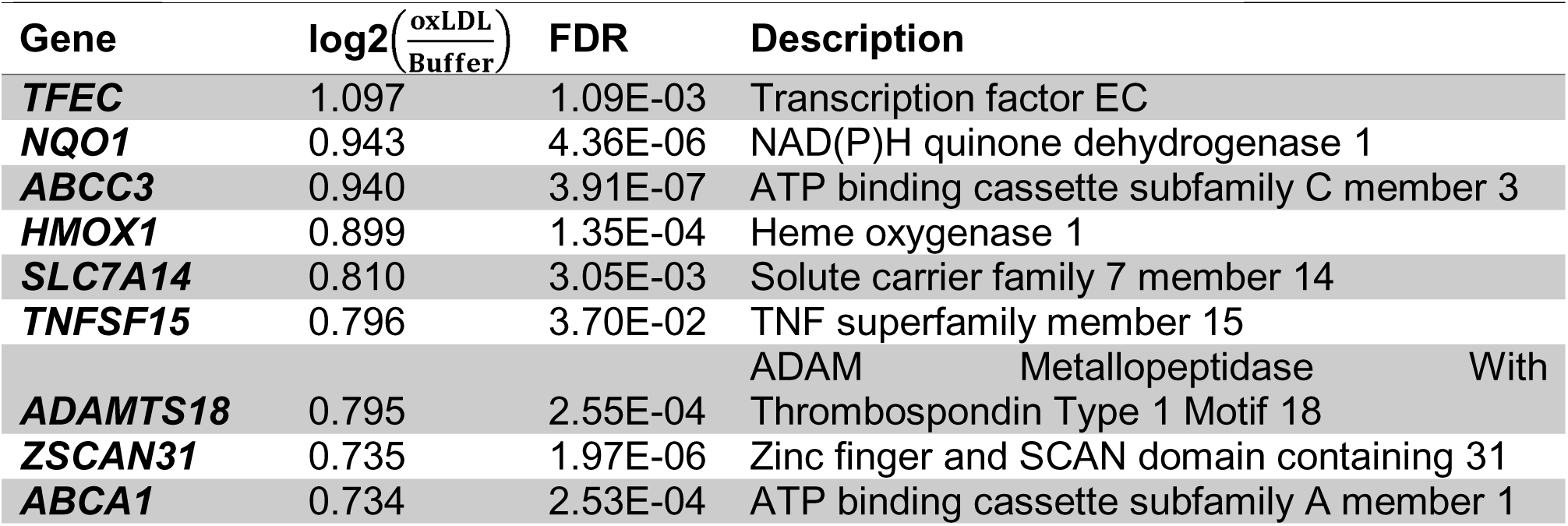
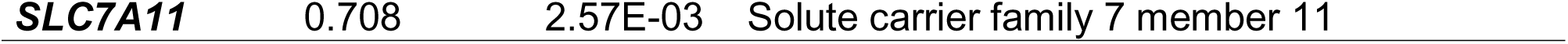
Top 10 most upregulated protein-coding genes by ox-LDL in HAECs.

**Table 2.** Top 10 most downregulated protein-coding genes by oxLDL in HAECs.

| Gene | $\log_2\left(\frac{\text{oxLDL}}{\text{Buffer}}\right)$ | FDR | Description |
| --- | --- | --- | --- |
| **CBLN2** | -0.936 | 3.33E-03 | Cerebellin 2 Precursor |
| **CCL23** | -0.932 | 1.55E-03 | Chemokine ligand 23 |
| **S1PR4** | -0.890 | 2.21E-02 | Sphingosine-1-phosphate receptor 4 |
| **MT1M** | -0.847 | 2.57E-03 | Metallothionein 1E |
| **C6orf132** | -0.800 | 4.67E-02 | Chromosome 6 Open Reading Frame 132 |
| **TXNIP** | -0.776 | 1.16E-02 | Thioredoxin-interacting protein |
| **DHCR24** | -0.765 | 3.91E-04 | 24-dehydrocholesterol reductase |
| **BDKRB2** | -0.729 | 3.34E-02 | Bradykinin Receptor B2 |
| **CD14** | -0.724 | 3.08E-02 | CD14 molecule |
| **COL21A1** | -0.717 | 3.01E-03 | Collagen Type XXI Alpha 1 Chain |

To investigate the biological pathways affected by ox-LDL, we performed an enrichment test for MSigDB and KEGG pathways (Figure 2B and Figure S3A). Consistent with our observation in macrophages, ox-LDL treatment triggered the anti-oxidative-stress NFE2 signalling pathway and inhibited intrinsic cholesterol biosynthesis (Figure 2B). Interestingly, we found that ferroptosis, an iron-dependent form of necrosis associated with endothelial dysfunction and increased atherosclerotic burden ^36^, was one of the most up-regulated pathways following ox-LDL exposure (Figure 2B and Figure S3A). This was observed even in the absence of any pro-adhesive, activated endothelial phenotypes.

The most upregulated protein-coding gene is *TFEC*, a member of the MiT/TFE transcription factor family. The MiT/TFE family has been linked to lysosome biogenesis in cancer cells ^37^, and we have recently demonstrated ox-LDL-induced expression of *MITF* in macrophages ^32^. However, the cellular function of *TFEC* is poorly characterised, and no other MiT/TFE members were differentially expressed in ox-LDL-exposed endothelial cells.

### Exposure to ox-LDL but not ox-PAPC recapitulates the transcriptomic signatures associated with endothelial cells in human atherosclerosis

We next sought to confirm whether the transcriptomic changes induced by ox-LDL exposure *in vitro* are present in human atherosclerosis *in vivo*. Briefly, the transcriptomic profiles of individual endothelial cells from human carotid plaques or adjacent disease-free vasculatures were analysed and aggregated into pseudo-bulk populations for differential expression tests, and the plaque expression signatures were compared with the ox-LDL expression signatures using a Rank-Rank Hypergeometric Overlap test ^38^.

It is important to note that the plaque endothelial cells were collected from advanced lesions, thus they were exposed to different microenvironments compared to our ox-LDL model. Despite this, we found that 65 out of the 304 differentially expressed genes induced by ox-LDL were also dysregulated in plaque (p = 3.25×10^-16^, Fisher’s Exact Test). Specifically, we identified significant overlaps at both ends of the differential expression gene list (Figure 2C). This enrichment, however, was not evident in endothelial cells treated with ox-PAPC ^22^ (Figure 2D). Moreover, only limited overlaps were found between genes regulated by ox-LDL and those regulated by ox-PAPC (Figure S3B-C). Clearly, our *in vitro* ox-LDL-exposure model provides new information that better characterises the role of endothelial cells in human atherosclerosis.

### Integrative analysis highlights a shared ox-LDL-response network between endothelial cells and macrophages

We next sought to characterise the potential regulators that drive the endothelial response to ox-LDL. To map the potential cis-regulatory elements in HAECs, we performed high-throughput sequencing experiments with Assay for Transposase Accessible Chromatin (ATAC-seq) and chromatin immunoprecipitation (ChIP-seq) targeting the active enhancer marker H3K27 acetylation (H3K27ac). Surprisingly, although the chromatin accessibility at transcription start sites positively correlates with the gene expression (Figure S4A), no region was confidently identified as differentially accessible in the ox-LDL group compared to the control group (Figure S4B).

We reasoned that gene expression changes in the absence of chromatin accessibility change could result from the re-organisation of transcription factor (TF) binding in pre-existing open chromatin. To investigate the underlying regulators driving the ox-LDL-induced gene expression, we performed in-silico deletion analysis using Lisa ^39^, integrating both the RNA-seq and ATAC-seq data (Figure 3A). In brief, Lisa first transforms open chromatin signals (or enhancer signals) into a gene-wise regulatory potentials score, based on the distance to the gene body and promoters. TF binding events, inferred from ChIP-seq datasets, are then deleted *in silico* from the original open chromatin signals. TFs are identified as potential regulators if their in-silico deletions lead to larger changes in regulatory potentials in the differentially expressed genes, compared to random background genes.

**Figure 3.**
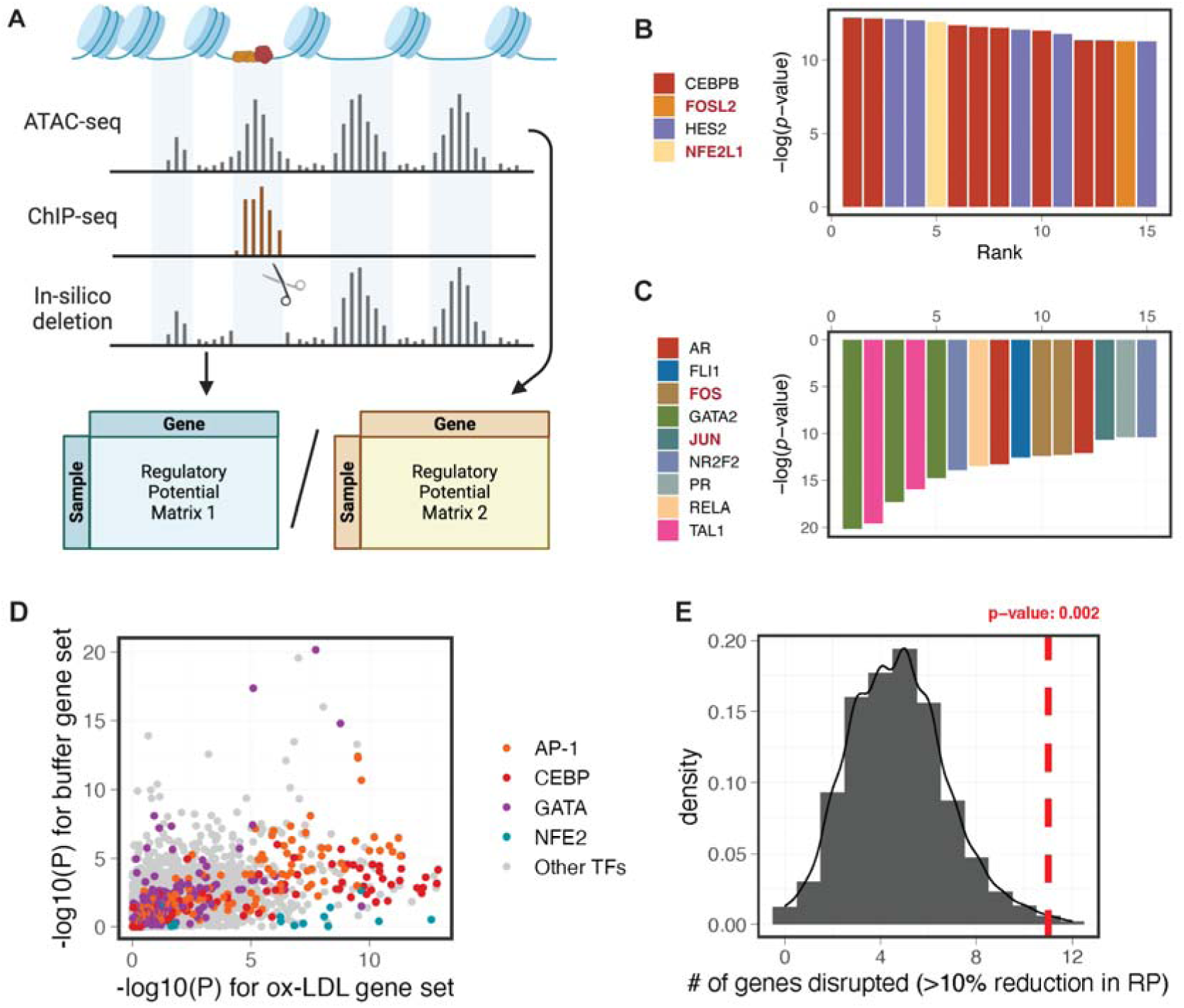
Integrative In-silico deletion (ISD) analysis identifies AP-1 and CEBP family factors as potential regulators underlying the transcriptomic response to ox-LDL. **A**, Overview of the in-silico deletion analysis: open chromatin signals were obtained from ATAC-seq data; for each transcription factor (TF), potential binding sites were inferred from public ChIP-seq data, and were computationally masked from the endothelial open chromatin landscape. The ATAC-seq signals before and after in-silico deletion were transformed into gene-wise regulatory potentials, and statistical significance was evaluated by comparing the changes in regulatory potential between differentially expressed genes and background genes. **B** and **C**, Candidate TFs regulating ox-LDL-induced genes (**B**) and ox-LDL-repressed genes (**C**) identified using the ISD analysis in LISA, ranked by significance. Each entry on the x axis represents an independent ChIP-seq experiment. **D**, ISD p-values of all expressed TFs (n=850 factors from n=5,587 ChIP-seq experiments) for ox-LDL and buffer gene sets. TFs in AP-1, CEBP, GATA and NFE2 families are highlighted. **E**, ISD of ox-LDL-altered CEBPB-binding sites significantly disrupts the regulatory potential of ox-LDL-induced genes as compared to expression-matched background genes. Genes with >10% reduction in regulatory potential after the in-silico deletion were considered as being disrupted. The empirical p-value was estimated from the null distribution using n=1000 expression-matched background genes.

Consistent with the pathway enrichment analysis (Figure 2B) and with our previous findings in macrophages ^32,40^, TFs from the AP-1 and NFE2 family were identified as the top regulators of both ox-LDL-induced genes (Figure 3B) and ox-LDL-repressed genes in HAECs (Figure 3C). We also repeated the in-silico deletion analysis with the H3K27ac ChIP-seq data and observed similar results (Figure S5A).

Our previous work has linked the transcription factor CEBPB to the cellular response to ox-LDL in human macrophages ^32^. In HAECs, CEBPB displayed the strongest ISD effect on ox-LDL-induced genes (Figure 3B and 3D), and was estimated to regulate about 20% of these genes (Figure S5B). The mRNA level of CEBPB, however, did not change after ox-LDL exposure in HAECs (Figure S5C).

To determine whether the regulatory effect of CEBPB is shared across different cell types, we next conducted an ox-LDL-specific in-silico deletion analysis. That is, instead of masking all CEBPB binding signals, we limited the in-silico deletion to regions where CEBPB binding was altered in ox-LDL-exposed macrophages, which represent only 8.7% of the total binding sites (8,718 out of 100,602 peaks, Figure S5D). Compared to randomly selected genes with similar expression levels, ox-LDL-induced genes were significantly more affected by this deletion of altered CEBPB binding sites (Figure 3E and Figure S5E). Collectively, our results demonstrate a shared ox-LDL-response network between endothelial cells and macrophages that is partly driven by the differential binding of CEBPB.

### Differential transcription factor bindings induced by ox-LDL activates endothelial cell migration

Our in-silico deletion analyses indicate that ox-LDL-induced transcriptomic changes are regulated by differential binding of transcription factors. To further characterise the genomic regions where these TFs act upon ox-LDL exposure, we carried out differential motif footprinting analysis using TOBIAS ^41^.

After correcting the Tn5 insertion bias, we identified 2,453 dynamic binding sites (DBSs) where the footprinting score for TF binding changed at least two-fold (n=1,172 and 1,281 sites for ox-LDL-activated and repressed regions, respectively). These DBSs were located across the genome (Figure S6A), and were enriched for H3K27ac signals compared to average open chromatin (Figure S6B).

Consistent with previous results, AP-1 and NFE-2 family factors were globally activated in ox-LDL-treated endothelial cells (Figure 4A), with an increase of 8% for AP-1 factor FOS (Figure S6C). CEBPB was not among the significant hits from footprinting analysis, possibly due to a known discrepancy between its *in vivo* binding pattern and the reported motif ^42^. Overall, these results suggest that ox-LDL treatment triggers a moderate change in TF binding within the already accessible chromatin.

**Figure 4.**
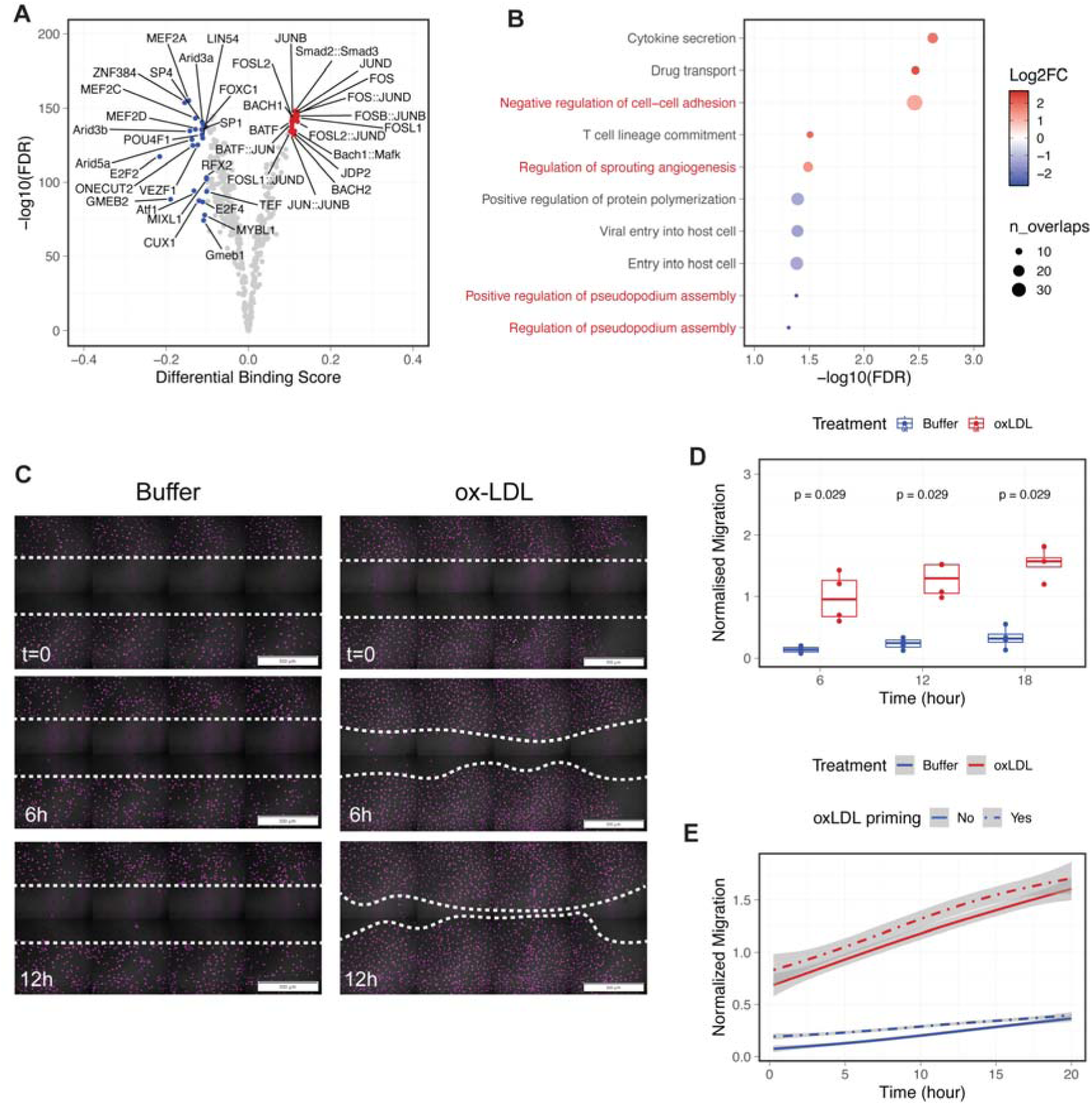
Changes in transcription factor binding activity indicate the activation of endothelial migration pathway in response to ox-LDL. **A**, Volcano plot of transcription factors with genome-wide binding patterns changed after ox-LDL exposure. Significant effect induced or repressed by ox-LDL (FDR<0.05, |Δscore| > 0.1) are coloured in red and blue, respectively. Only TF expressed in HAECs were included in this analysis. **B**, Predicted functions of the differential binding sites (DBS) using the Genomic Regions Enrichment of Annotations Tool (GREAT). Colour represents log2-fold change and size represents the number of overlapped genes in each annotation. Pathways associated with endothelial migration are highlighted in red. **C**, Representative live-cell fluorescence images of SiR-DNA labelled nuclei at specified time points, with a uniform gap introduced by culture inserts at the start of the assay. Scale bar: 500um. **D**, Endothelial migration quantified as the percentage of cells in the gap region normalised to that in buffer-treated cells (n=4 biological replicates). P-values calculated using Wilcoxon rank-sum tests. **E**, Normalised migration of HAECs with or without 24 hours of ox-LDL priming. Shaded regions indicate 95% confidence intervals estimated from n=4 biological replicates.

We next performed a pathway enrichment test in DBSs to explore the biological consequence of the observed change in TF binding. Notably, functional annotations related to endothelial migration were highly enriched in genes near both the ox-LDL-activated and ox-LDL-repressed DBSs (Figure 4B). To experimentally validate this finding, we further performed cell migration assays, and found that ox-LDL exposure in HAECs significantly increased endothelial cell migration (Figure 4C and 4D). This experiment was also repeated with cells pre-treated with ox-LDL, and the effect of ox-LDL priming only lasted a few hours after the start of the assay, with no significant difference after 20 hours (Figure 4E). This observation echoes previous results where we detected no global alteration in chromatin accessibility, suggesting a fast and direct endothelial response to ox-LDL exposure.

Together, our computational and experimental results demonstrate that ox-LDL triggers a rapid activation of endothelial migration, potentially mediated by genome-wide epigenomic changes in transcription factor binding.

### Prioritisation of CAD variants using the DBSs in endothelial cells

Endothelial cells are one of the most important arterial cell types that are enriched for the disease heritability of CAD ^27,28^. Given that ox-LDL triggers major changes in metabolism, transcriptome and epigenome in HAECs, we hypothesised that the endothelial response to ox-LDL is also genetically linked to human atherosclerosis.

To test this hypothesis, we performed an enrichment test for CAD heritability in dynamic TF binding sites. Similar to previous reports, endothelial open chromatin regions that contain all the TF binding sites are enriched for the SNP heritability of CAD (Figure 5A). This enrichment was not observed for irrelevant neurological traits like smoking behaviour. In addition, we found that the ox-LDL-induced differential binding sites carry 4-times higher per-SNP heritability compared to average binding sites (Figure 5A), indicating that the genetic risk of CAD is concentrated in the TF binding sites altered by ox-LDL exposure.

**Figure 5.**
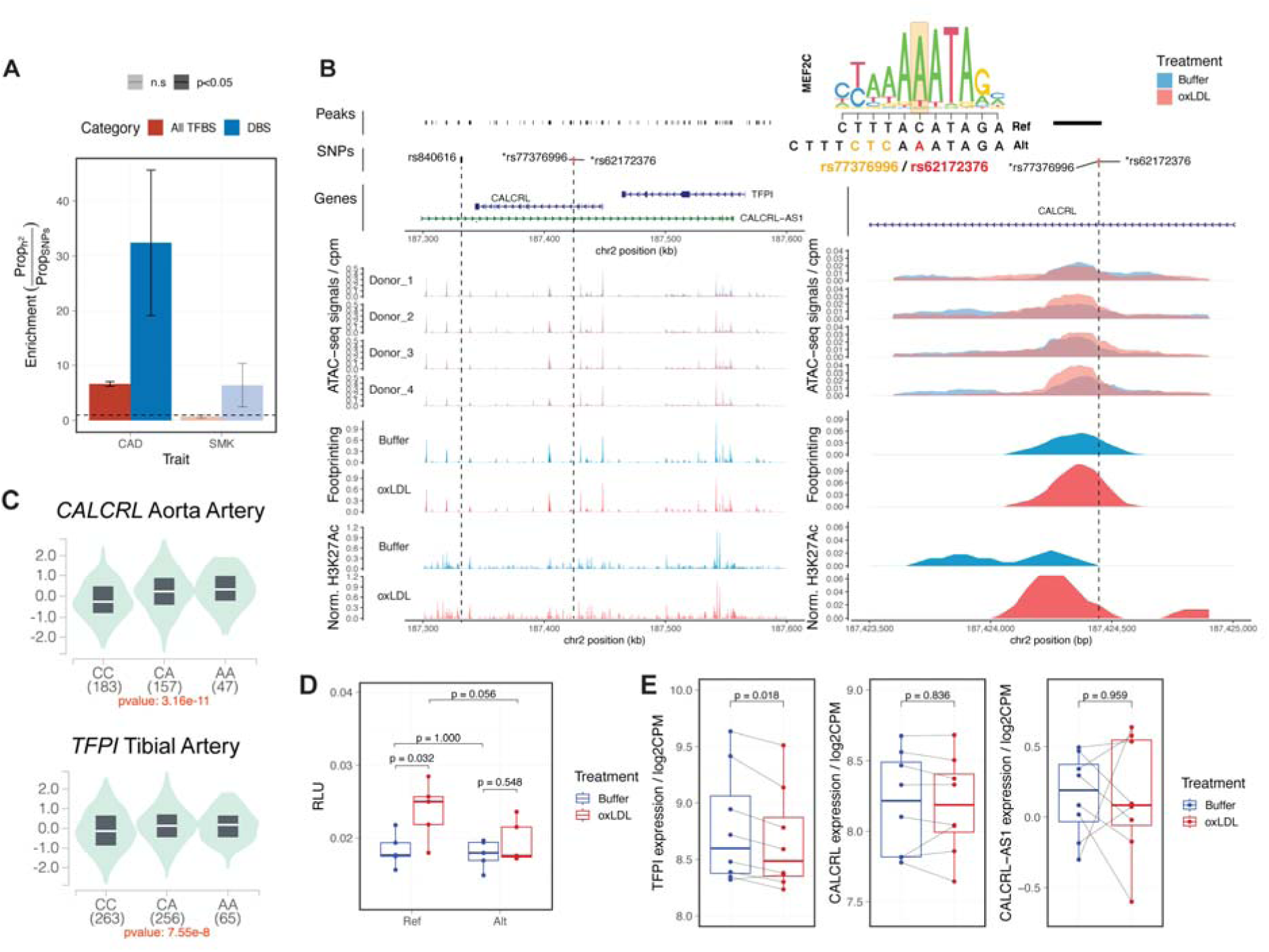
Integrative analysis links CAD risk variants rs62172376 to *TFPI* and reveals allelic-specific activation induced by ox-LDL. **A**, Enrichment test for disease heritability in open chromatin regions containing foot prints from TF binding (All TFBS) and dynamic TF binding sites (DBS). Standard errors for enrichment statistics were estimated using s-LDSC. Non-significant enrichment results are plotted in transparent. Enrichment for heritability of smoking behaviour (SMK) is shown to the right as a negative control. GWAS summary statistics were obtained from GWAS Catalog using study ID GCST005194 ^24^, and GCST007327 ^46^ for CAD and SMK, respectively. **B**, The regulatory landscape at the prioritised *CALCRL*/*TFPI* locus. The tagged variant rs840616 is marked in grey and two candidate causal variants rs77376996 and rs62172376 are marked in red. Tracks (from top to bottom) showing: ATAC-seq peaks (peaks), CAD-associated variants (SNPs), genomic annotations (Genes), ATAC-seq signals aggregated by biological donors, Motif footprinting score, and enhancer signals from H3K27ac ChIP-seq aggregated by treatment. Higher resolution coverage plots centred on the risk variant are plotted in the right panel. **C**, Violin plot showing the allele-specific effect of the haplotype on the expression of *CALCRL* and *TFPI* in human artery based on the GTEx data. Samples are grouped by the genotype at the SNP rs62172376. **D**, Dual Luciferase reporter assay of the 150bp dsDNA construct containing each haplotype. HAECs were transfected with the indicated plasmid, treated with or without ox-LDL for 24 hours and lysed for luciferase activity measurement (n=5 replicates per group). Signals normalised to the Renilla luciferase activity and reported as relative light units (RLU). P-values calculated with Wilcoxon tests. **E**, Normalised gene expression (log2 count-per-million-reads) of *TFPI*, *CALCRL* and *CALCRL-AS1* in unexposed and ox-LDL-exposed HAECs (n=4 biological replicates with 2 technical repeats per donor). P-values were calculated using the quasi-likelihood test in edgeR with a linear mixed model controlling both the donor and batch effect.

Of the 330 independent risk loci that reach the genome wide significance threshold (p<5×10^-8^) in genome-wide association studies (GWAS) ^25,26^, 5 contain variants that reside in dynamic binding sites (Table 3). The risk locus at chromosome 9 (9p21.3) has been extensively characterised using knockout mouse models ^43^, although no eQTL signals were found at the locus in human endothelial cells and vascular smooth muscle cells.

**Table 3.** Prioritised CAD-risk variants in ox-LDL-induced DBSs.

| Rsid | Chr | Pos | Ref | Alt | DBS | Annotation |
| --- | --- | --- | --- | --- | --- | --- |
| rs56170783 | chr1 | 56550459 | A | C | chr1-56549815-56550830 | PLPP3 intron |
| **rs17048367** | chr1 | 218660548 | A | G,T | chr1-218660506-218661540 | Intergenic region |
| **rs7068966** | chr10 | 12235993 | C | G,T | chr10-12234966-12236001 | CDC123 intron |
| **rs77376996** | chr2 | 187424445 | T | TCTC, TCTCA | chr2-187423908-187424925 | CALCRL intron |
| **rs62172376** | chr2 | 187424447 | C | A | chr2-187423908-187424925 | CALCRL intron |
| **rs1537372** | chr9 | 22103184 | G | A,T | chr9-22103132-22104146 | CDKN2B intron |
| **rs1537373** | chr9 | 22103342 | T | A,G | chr9-22103132-22104146 | CDKN2B intron |
| **rs1333042** | chr9 | 22103814 | A | C,G | chr9-22103132-22104146 | CDKN2B intron |

The *PLPP3* gene has been linked to endothelial barrier function in various experimental models ^44,45^, and a SNP at the *PLPP3* locus was found in an endothelial-specific enhancer reported in a previous variant prioritisation study ^22^. Here, we found that another risk variant rs56170783 at this locus overlaps an ox-LDL-induced TF binding site, and displays allelic imbalance in ATAC-seq signals from the heterozygous donor (Figure S7A and B).

### Prioritised variants at the *CALCRL*/*TFPI* locus

At the *CALCRL*/*TFPI* locus, a genetic variant rs840616 has been associated with CAD risks (β=0.0402) in European populations ^24^. In linkage disequilibrium with this lead variant (r^2^=0.81), two closely-linked variants rs77376996 and rs62172376 overlapped with an intronic DBS in the *CALCRL* gene (Figure 5B). Although the increase in chromatin accessibility at this site was not significant after FDR correction, our motif footprinting analysis indicates a more than two-fold increase in TF binding activity (Figure 5B). This ox-LDL-induced activation is further supported by the substantial increase in H3K27ac signals in adjacent histones (Figure 5B).

There are two protein-coding genes *CALCRL* and *TFPI* expressed at this locus, whose transcription start sites are 24kb and 141 kb downstream of the SNP rs62172376, respectively. This locus also contains a long non-coding RNA *CALCRL-AS1* that is antisense to *CALCRL* and *TFPI*. The protective haplotype (rs62172376-A) contains a MEF2 binding site (Figure 5B) and in the GTEX database is linked to higher expression of both genes *CALCRL* and *TFPI* in human artery (Figure 5C).

The allele-specific activation in ox-LDL-exposed endothelial cells was confirmed using dual luciferase reporter assays, where the regulatory element containing the risk reference haplotype displays higher enhancer activity after ox-LDL exposure (Figure 5D). Interestingly, despite being closer to the DBS, *CALCRL* did not exhibit an ox-LDL-induced change in expression (Figure 5E). In fact, *TFPI* is the only gene at this locus that was differentially regulated by ox-LDL (log2FC=-0.12, p=0.018; Figure 5E). These results indicate that *TFPI* but not *CALCRL* is more likely to be the potential causal gene at the CAD risk locus containing rs62172376 which may protect endothelial cells from ox-LDL.

Of note, a closer inspection of molecular QTL signals highlights an alternative splicing event of the anti-sense non-coding RNA *CALCRL-AS1* associated with rs62172376 in human coronary artery (Figure S8A and B). Specifically, we found that the last intron of *CALCRL-AS1* was partially retained in endothelial samples carrying the protective haplotype (Figure S8A). This intron is anti-sense to the promoter sequence of *TFPI*, and its usage positively correlated with the expression of *TFPI* (Figure S8C), but not *CALCRL* (Figure S8D) in HAECs.

Together, our integrative analyses pinpoint a causal variant rs62172376 at the *CALCRL*/*TFPI* locus, prioritise the gene *TFPI* as the target gene, and capture a gene-environment interaction underlying the cellular response to ox-LDL in endothelial cells.

## Discussion

The initial exposure of endothelial cells to ox-LDL marks one of the earliest pathological events in atherosclerosis. In this study, we offer the first comprehensive transcriptomic and epigenomic characterisation of the response to ox-LDL in primary human aortic endothelial cells. Our findings reveal that ox-LDL triggers changes in the transcriptome and cellular metabolism that are non-inflammatory. By integrating multi-omics sequencing data, we are able to identify the TFs that drive the ox-LDL response, which features a transcriptional regulatory network that is largely shared with human macrophages. The redistribution of these TFs also activates endothelial-specific pathways, including migration. Crucially, we demonstrate that changes in TF binding induced by ox-LDL are enriched for CAD heritability, and can be utilised as a leverage point to prioritise causal risk variants involved in the endothelial response to atherogenic lipids.

In HAECs, exposure to ox-LDL reduces mitochondrial respiration, thereby shifting the energy balance towards glycolysis. This effect is contradictory to the effect of ox-PAPC, a widely studied synthesised bioactive component of ox-LDL, which has been reported to increase the respiration and metabolic activity in HAECs ^23^. *In vivo*, we demonstrated a robust correlation between human atherosclerotic lesions and the effect of ox-LDL, but not for ox-PAPC. Clearly, the proatherogenic effect of ox-LDL is a combination of complex, sometimes opposing influences from individual components. Studying one single element within the ox-LDL mixture, despite the increased biochemical specificity and the ease of preparation, may miss the whole picture and so could lead to misleading conclusions.

With a comprehensive genome-wide footprinting analysis, we discovered an activating effect of ox-LDL on endothelial migration. This finding was functionally validated using live-cell microscopy. It is worth noting that contrasting phenotypes have been reported in different endothelial lines at ox-LDL concentrations ranging from 1 to 50 ug/mL ^47–49^. In HUVECs, ox-LDL stimulates migration at 5 ug/mL, but inhibits it at 40 ug/mL ^47^. On a molecular level, the impaired migration capacity was linked to the dephosphorylation of Akt in HUVECs ^48^. However, increased Akt phosphorylation has been reported in HAECs after ox-LDL treatment ^50^. Given that HUVECs are venous cells derived from immune-privileged foetal tissue, they may be less tolerant to the oxidative stress compared to adult arterial endothelium. This could also account for the absence of endothelial activation phenotypes in our *in vitro* model using aortic endothelial cells. Nevertheless, we show that the non-inflammatory response to ox-LDL, which drives the migration phenotype, is significantly enriched for disease heritability. In other words, this supports the ox-LDL-induced *in vitro* migration phenotype being causally involved in the *in vivo* progression of human atherosclerosis.

Since many non-coding genetic variants influence disease risks by altering the function of regulatory DNA ^51^, profiling the colocalization of regulatory regions with genetic risk variants in LD offers a powerful approach to decipher the causal variants and their target genes ^52^. The prioritised risk locus around the *CALCRL* and *TFPI* gene is particularly interesting, as only one of the two coding genes is differentially expressed by ox-LDL exposure. The dynamic binding site containing the risk reference haplotype displayed significant activation after ox-LDL treatment, while the protective haplotype remained unchanged. This change in enhancer activity was not found under basal conditions, which may explain why rs62172376 was missed in previous variant prioritisation studies. In fact, our results illustrate a context-specific regulatory effect at this risk locus, underscoring the necessity to adapt experimental models with disease-relevant stimuli in the functional fine-mapping of GWAS results.

Overall, this study provides a comprehensive understanding of the transcriptional regulation of ox-LDL response in human endothelial cells. Our multi-omics data lays a solid foundation for future research exploring the interactions between the genetic and environmental component of CAD.

## Materials and Methods

### Data Availability

The raw sequencing data has been deposited in the National Center for Biotechnology Information Sequence Read Archive (NCBI-SRA) under accession number PRJNA1145858. A temporary reviewer link has been created to access the metadata: https://dataview.ncbi.nlm.nih.gov/object/PRJNA1145858?reviewer=r8gnl22cqqbr0dmo1l3b51nq0b. Raw and processed data will be released upon the publication of this manuscript.

### Cell Culture

Primary human aortic endothelial cells (HAECs) were obtained from Promocell (Promocell, Heidelberg, Germany) and expanded at passage 2 before usage. HAECs were maintained in Endothelial Cell Growth Medium MV2 (Promocell, Heidelberg, Germany) supplemented with 100 units/mL penicillin and 100 ug/mL streptomycin. Cells were assayed at ∼80% confluency between passage 4 to passage 7. Due to the privacy policy, no demographic information other than the catalogue number of donors can be retrieved from the company. The catalogue numbers for the four HAEC lines used in this study are: #4082102.16, #422Z037, #431Z002.3, #452Z031.1.

### Cell Treatments

LDL was purified from human plasma by isopycnic ultracentrifugation, then oxidized overnight using 25 uM CuCl_2_, as described previously ^40^. DPBS solution (Thermo Scientific, Waltham, MA) from the last round of dialysis was used as the control buffer. The concentration of ox-LDL was determined using the BCA protein assay (Thermo Scientific). Unless otherwise specified, cells were treated with a final concentration of 50 ug/mL of ox-LDL or an equal volume of control buffer for 48 hours. For immunofluorescence studies, LDL was labelled overnight at 37 C with 300 ug DiI per mg LDL, before purification by ultracentrifugation and oxidization using CuCl_2_.

### Flow Cytometry

Before the assay, HAECs were treated with 50 ug/mL of DiI-labelled ox-LDL or control buffer, or 10 ug/mL recombinant human TNF alpha protein (ab9642, Abcam) for 24 hours. Cells were detached using StemPro Accutase solution (Thermo Scientific) and re-suspended in ice-cold FACS buffer (PBS, 1% BSA and optional 1% azide). Primary antibody staining was carried out in the dark at 4 C for 30min with the following antibody dilutions in FACS buffer: 1:200 ICAM-1 (353102, Biolegend), 1:500 VCAM-1 (14-1069-82, Invitrogen). Second staining was performed with 1:1000 Anti-Mouse-IgG-647 (A-21240, Invitrogen) at 4 C for 30min. Zombie Violet (423113, Biolegend) was included as a viability dye in a final dilution of 1:200. Data was exported to FlowJo 10 (FlowJo LLC, Ashland, USA) for quantitative analysis.

### Immunofluorescence Imaging

For fixed cell imaging, HAECs were seeded at 15,000 cells/well in 8-well ibiTreat μ-slides (Ibidi) and assayed at ∼80% confluency before treatment with 50 ug/mL DiI-labelled ox-LDL or control buffer. Cells were fixed at room temperature for 15 mins with 4% PFA in PBS, followed by permeabilization with 0.1% Triton X-100 (Thermo Scientific) and 0.2% Tween-20 (Roche) in PBS. Fixed cells were incubated in blocking buffer (1% BSA, 2% FCS, 0.3M glycine and 0.1% Tween-20 in PBS) at room temperature for 1 hour. Cytoskeleton was staining with 1:500 tubulin-AF488 (clone DM1A, eBioscience) and nuclei were counterstained with Hoechst 33342 (Santa Cruz Biotechnology) diluted 1:10,000 in PBS. Stained cells were imaged on a LSM 900 with Airyscan2 in confocal mode (Zeiss).

For the migration assay, cell culture plates with 2-well silicone inserts (80241, Ibidi) were used to create a 500 um cell-free gap for uniform imaging of endothelial migration. Prior to the assay, HAECs were seeded at the density of 10,000 cells per culture insert well and were starved in supplement-depleted EC culture medium containing 50ug/mL ox-LDL or control buffer overnight. Before the assay, cells were pre-stained with 300 nM SiR-DNA (CY-SC007, Universal Biologicals) for at least 2 hours. Culture inserts were carefully lifted using a sterile tweezer, and cells were gently washed with sterile PBS twice. Live cells were then imaged every 15 minutes for 24 hours in assay media containing 300 nM SiR-DNA and 50 ug/mL ox-LDL or control buffer on the Olympus SpinSR SoRa and ScanR High Content system (Olympus). The location of nuclei was recovered using arivis Vision4D version 4.1 (Zeiss), and the migration rate was defined as the percentage of nuclei overlapping the initial gap region. Migration rates were normalised to the buffer group within paired samples to mitigate any donor effect.

### Seahorse

Metabolic measurements of HAEC were performed with Seahorse XF Mito Stress Test and XF Glycolytic Rate Assay kits using a Seahorse XFe96 Analyzer (Agilent Technologies, Santa Clara, US). 50,000 endothelial cells were seeded into the Seahorse 96-well plates and were treated with 50ug/mL oxLDL or control buffer 24 hours prior to the assay. For mito stress test, 2uM of Oligomycin, 1uM of FCCP and 0.5uM of Rotenone/Antimycin A were used based on a pilot titration experiment; For glycolytic rate, 0.5uM of Rotenone/Antimycin A and 50mM of 2-DG were used. Results were analysed using Seahorse Wave Desktop (Agilent).

### Library preparation for high-throughput sequencing

Total RNA from HAECs was purified using RNeasy Micro Kit (Qiagen) after TRIzol and chloroform extraction according to the manufacturer’s protocol. The quality of RNA samples was analysed using a TapeStation 2200 with High Sensitivity RNA ScreenTape (Agilent Technologies, Santa Clara, US). Samples with high integrity (RIN score>9) were selected for PolyA-enrichment and TruSeq library preparation (Illumina). Each sample was sequenced to a target depth of 37.5 million read pairs using a NovaSeq 6000 device (Illumina).

ATAC-seq libraries were prepared according to the Omni-ATAC protocol ^53^. In brief, nuclei were isolated by incubation with ATAC-Resuspension Buffer (ATAC-RSB) containing 0.1% NP40 (Roche, Basel, Switzerland), 0.1% Tween-20 (Roche, Basel, Switzerland) and 0.01% Digitonin (Promega, Madison, WI) on ice for 3 minutes, and were washed with 1 mL of ATAC-RSB containing 0.1% Tween-20. Tn5 tagmentation reaction was performed at 37°C for 30 minutes using the Nextera kit (Illumina). ATAC-seq libraries were then PCR amplified using NEBNext High-Fidelity PCR master mix (New England Biolabs, Ipswich, US) and index primers from Nextera Kit (Illumina), and analysed using a TapeStation 2200 with High Sensitivity D1000 ScreenTape (Agilent). Libraries showing successful transposition were sequenced to a target depth of 112.5 million read pairs using a NovaSeq 6000 device (Illumina).

ChIP-seq was performed using the MAGnify Chromatin Immunoprecipitation System (ThermoFisher Scientific) following the manufacturer’s instructions. HAECs were resuspended in 1% formaldehyde at room temperature for 10 minutes, and the cross-linked chromatins were fragmented using a Bioruptor (Diagenode) with 16 cycles of 30 seconds on and off. 500,000 cells per IP were incubated with 100uL Dynabeads conjugated with 2 ul of Anti-H3K27ac (ab472, Abcam) or Anti-Rabbit-IgG control (Thermo Scientific) at 4°C overnight. After reverse crosslinking, ChIP-ed DNA was purified using MinElute Kit (Qiagen) followed by TruSeq library preparation (Illumina). were sequenced to a target depth of 40 million read pairs using a Hi-Seq 2500 device (Illumina).

For all bulk sequencing libraries, read qualities were checked using FastQC ^54^. When adapter contamination was detected, the 3’ end of affected reads was trimmed using NGMerge ^55^.

### RNA-seq Data Analysis

Adaptor-trimmed reads were first aligned to the reference human genome using the splice-aware STAR aligner ^56^. Human genome assembly GRCh38.p13 and gene annotation Release 43 were downloaded from the GENCODE site and were indexed using STAR. SAMtools ^57^ was used to filter out unmapped or multi-mapping reads, and gene-wise count matrices were generated using featureCounts of the Subread package^58^. Differential expression analysis was performed in R using edgeR ^59^. Genes with low expression (min.count<50 in more than half of the samples) were filtered out, and the remaining count matrices were normalised by sequencing depth and by the Trimmed Mean of M-values (TMM) method. Differential gene expression between treatment groups was tested using the quasi-likelihood test, with a generalized linear model controlling for experimental batches and biological donors. The test statistics were adjusted using the Benjamini-Hochberg (B-H) method and genes with adjusted p-values <0.05 were considered as differentially expressed. Pathway enrichment analyses were performed using the XGR package ^60^, with functional annotations collected from MsigDB and KEGG.

Single-cell RNA-seq profiles of human carotid atherosclerotic plaque and adjacent control tissue were retrieved from Gene Expression Omnibus using the accession number GSE159677 ^61^, as part of our recent meta-analysis in ^32^. Endothelial cells were selected based on gene expression of *PECAM1*, *VWF* and *PLVAP*, and were aggregated into pseudo-bulk population per sample. Differential expression tests were performed using edgeR with the quasi-likelihood test, with donor information included as a covariate. The Rank-Rank Hypergeometric Overlap tested were conducted in R using the package RRHO ^38^, and genes were ranked by sign(log-fold-change)xlog_10_(P).

### ATAC-seq Data Analysis

Adaptor-trimmed reads were aligned to the hg38 reference genome using Bowtie2 with --very-sensitive settings ^62^. PCR duplicates were marked using Picard, and were further removed from downstream analysis together with mitochondrial reads using SAMtools. Post-alignment QC was performed as instructed by the ENCODE ATAC-seq processing pipeline, and all libraries passed the threshold of Non-Redundant Fraction>0.9, PCR Bottleneck Coefficients-1>0.9, PCR Bottleneck Coefficients-2>3, and Transcription Start Site Enrichment score >7.

For downstream analysis, reads were trimmed using BEDTools ^63^ to retain only the 9-bp Tn5 cutting sites at both ends of the DNA fragments. A consensus peak set was called using MACS3 ^64^ combining all sequenced samples, with peaks overlapping the ENCODE blacklisted region being removed before read counting in Subread. Differential accessibility analysis was performed using edgeR, as described above. A window-based differential accessibility test was also performed by counting reads within 100bp sliding windows across the genome using csaw ^65^.

To determine the donor genotype, aligned ATAC-seq reads were aggregated by sample and genotyped using the GATK (v4.1.7.0) germline short variant discovery pipeline ^66^. PCR duplicates were marked with MarkDuplicates. Base quality scores were recalibrated against dbSNP155 known sites, and variants were called with HaplotypeCaller, combined using GenomicsDBImport, and genotyped with GenotypeGVCFs. Genotyping results from SNPs in perfect LD (r^2^=1 in 1000G EUR samples) were merged.

### ChIP-seq Data Analysis

As described above, adaptor-trimmed reads were aligned to the hg38 reference genome using Bowtie2 followed by PCR duplicates removal using Picard. For H3K27ac ChIP-seq, aligned bam files were merged with subsampling per treatment group. Aggregated ChIP-seq signals were computed over 100bp bins and normalized by sequencing depth using the bamCoverage from DeepTools, with 150bp extension of each fragment. For function annotation of dynamic binding sites (DBSs), ChIP-seq reads were counted per 100bp bin for each 4kb window centred at each DBS, normalized by sequencing depth, and smoothed using a generalized additive model implemented through ggplot2. The macrophage CEBPB ChIP-seq dataset was generated previously and retrieved from GEO (GSE54975) ^40^. A consensus peak set was called using MACS3 with all samples, and peaks overlapping the ENCODE blacklisted region were removed. Differential binding tests were performed in edgeR, with both donor and treatment included as covariates in the quasi-likelihood test. Peaks with p-values lower than 0.05 were retained (n=8,718) for ox-LDL specific in-silico deletion analysis.

### In-Silico Deletion Analysis

In-silico deletion (ISD) tests were performed using the “FromCoverage” function of the Lisa package ^39^ with 500 background genes and the default ‘enhanced_10k’ regulatory potential model. The consensus open chromatin profile from ATAC-seq or the consensus enhancer profile from H3K27ac ChIP-seq was used to compute the background regulatory potential. For the ISD of ox-LDL-specific CEBPB bindings, fragments overlapping altered CEBPB binding sites were removed using BEDTools, and the gene-wise regulatory potentials were calculated using the “genescore” function implemented through the MAESTRO package ^67^; empirical null distribution was estimated from n=1000 sets of expression-matched background genes.

### Differential Motif Footprinting Analysis

TOBIAS ^41^ was used to analyse the foot print signatures in ATAC-seq data. In brief, the merged 9-bp consensus bam file was processed using ATACorrect function in TOBIAS to correct for the intrinsic bias in Tn5-cut sites; Footprinting scores were calculated at base-pair resolution within open chromatin peaks using ScoreBigwig; differential foot printing was then conducted using BINDetect, with transcription factor (TF) motifs downloaded from JASPAR core 2018 ^68^. For ontology enrichment and heritability analysis, differential TF binding sites was extended 500bp at each end; the enrichment for functional annotations was tested using GREAT v4 ^69^.

### GWAS and QTL SNPs

GWAS statistics were curated by NHGRI-EBI Catalog and downloaded from the UCSC hg38 database (April 2022). We identified SNPs with significant genome-wide association (p<5×10^-8^), pruned the list (r^2^ > 0.1 identified by 1000 Genome Project phase 3 in EUR samples, hg38 build) to remove redundant loci, and then pooled all SNPs in high LD within each locus (r^2^ > 0.8 in EUR samples) to establish a set of trait-associated variants. Only dbSNP common variants (MAF>1%, dbSNP155) were considered in this study. LD information was extracted using PLINK v1.90b6.26 ^70^. eQTL SNPs were obtained from GTEx v8 (Aorta Artery, Coronary Artery and Tibial Artery) ^71^ at hg19 coordinates were converted to hg38 using annotations from NCBI dbSNP Build 155.

### Heritability Analysis

LDSC v1.01 was used to partition CAD heritability and test the enrichment for DBSs as described ^72,73^. The standard errors of enrichment statistics and regression coefficients are computed using a block jack-knife (n=1,000) ^72^. The full summary statistics for CAD used in this study were downloaded from the GWAS Catalog, study ID GCST005194 ^24^. Smoking behaviour was included as a negative control and its summary statistics were retrieved using with the ID GCST007327 ^46^.

### Dual Luciferase Assay

The 150-bp enhancer element containing either the reference or the alternative haplotype centred at rs62172376 was synthesized (GenScript Biotech, Piscataway, US) and cloned into empty pGL4.27 Firefly luciferase reporter (Promega, Fitchburg, WI). Successful cloning was verified by sequencing prior to transfection. For dual luciferase assay, 1 ug pGL4.27 Firefly luciferase reporter plasmid and 0.02 ug pRL-SV40 Renilla luciferase plasmid were co-transfected using ViaFect Transfection Reagent (Promega) in a 1:6 ratio per well in 12-well plates. HAECs were treated with 50 ug/mL ox-LDL or buffer 24 hours post transfection for another 24 hours, and were lysed for luciferase assay using the Dual Luciferase Reporter Assay System (Promega). Luciferase activities were measured using a CLARIOstar Microplate Reader (BMG Labtech, Ortenberg, Germany), and Firefly luciferase activity was normalized to Renilla luciferase activity as relative light units.

## Acknowledgments

The authors would like to thank the Oxford Genomic Centre and the Cellular Imaging Core at the Centre for Human Genetics for their support. We thank Dr Thomas Nicol from the Department of Physiology, Anatomy & Genetics, University of Oxford for the help with metabolic assays. We also thank Prof. Julian Knight at the Centre for Human Genetics and Dr Mark Crabtree at the University of Surrey for their insightful comments during the preparation of this manuscript. Illustrations in Figure 3A were created with BioRender.com.

## Author Contributions

Study conception and design: J.J. and C.A.O. Acquisition of data: J.J., T.K.H, A.C., Y.M. and T.A. Analysis and interpretation of data: J.J. and C.A.O. Drafting and writing of manuscript: J.J. and C.A.O.

## Supplementary Materials

**Figure S1.**
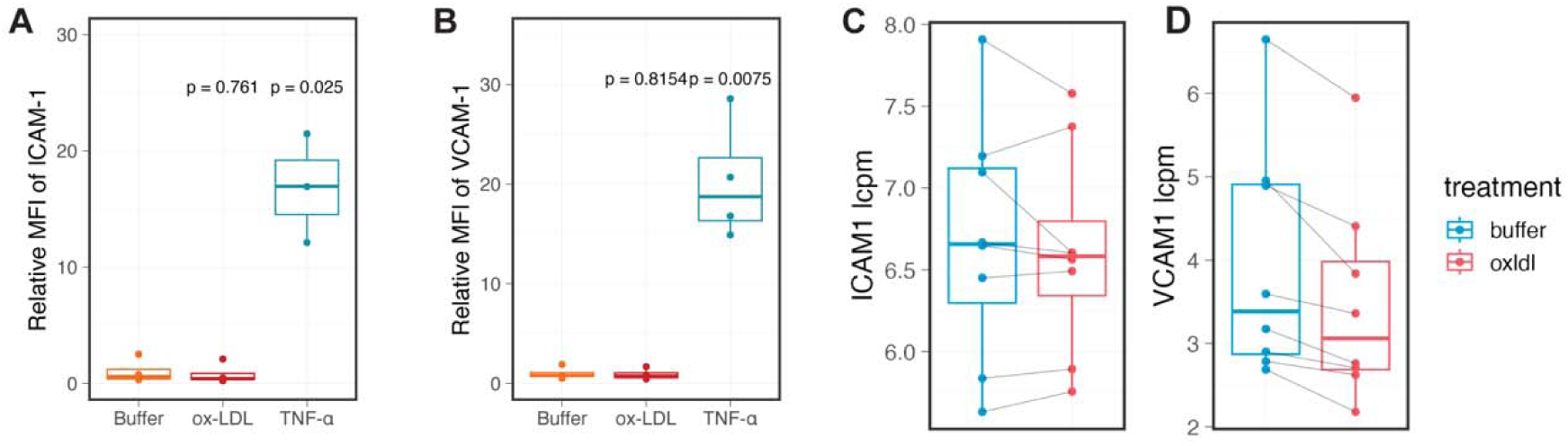
Ox-LDL alone does not lead to a typical activated endothelial phenotype. **A** and **B**, Quantification of membrane ICAM-1 (**A**) and VCAM-1 (**B**) level using flow cytometry in HAECs treated with 50ug/mL ox-LDL or 10ng/mL TNF-α for 24 hours (n=4 biological replicates). Signals were normalised to the buffer group and shown as the relative mean fluorescence intensity. P-values were calculated using t-tests. **C** and **D**, Normalised log-count-per-million gene expression of *ICAM1* (**C**) and *VCAM1* (**D**) as quantified by bulk RNA-seq, coloured by treatment group. HAECs were treated with 50ug/mL ox-LDL for 48 hours for RNA-seq experiments (n=4 biological replicates with 2 technical repeats per donor). Paired samples from the same donor are connected in lines.

**Figure S2.**
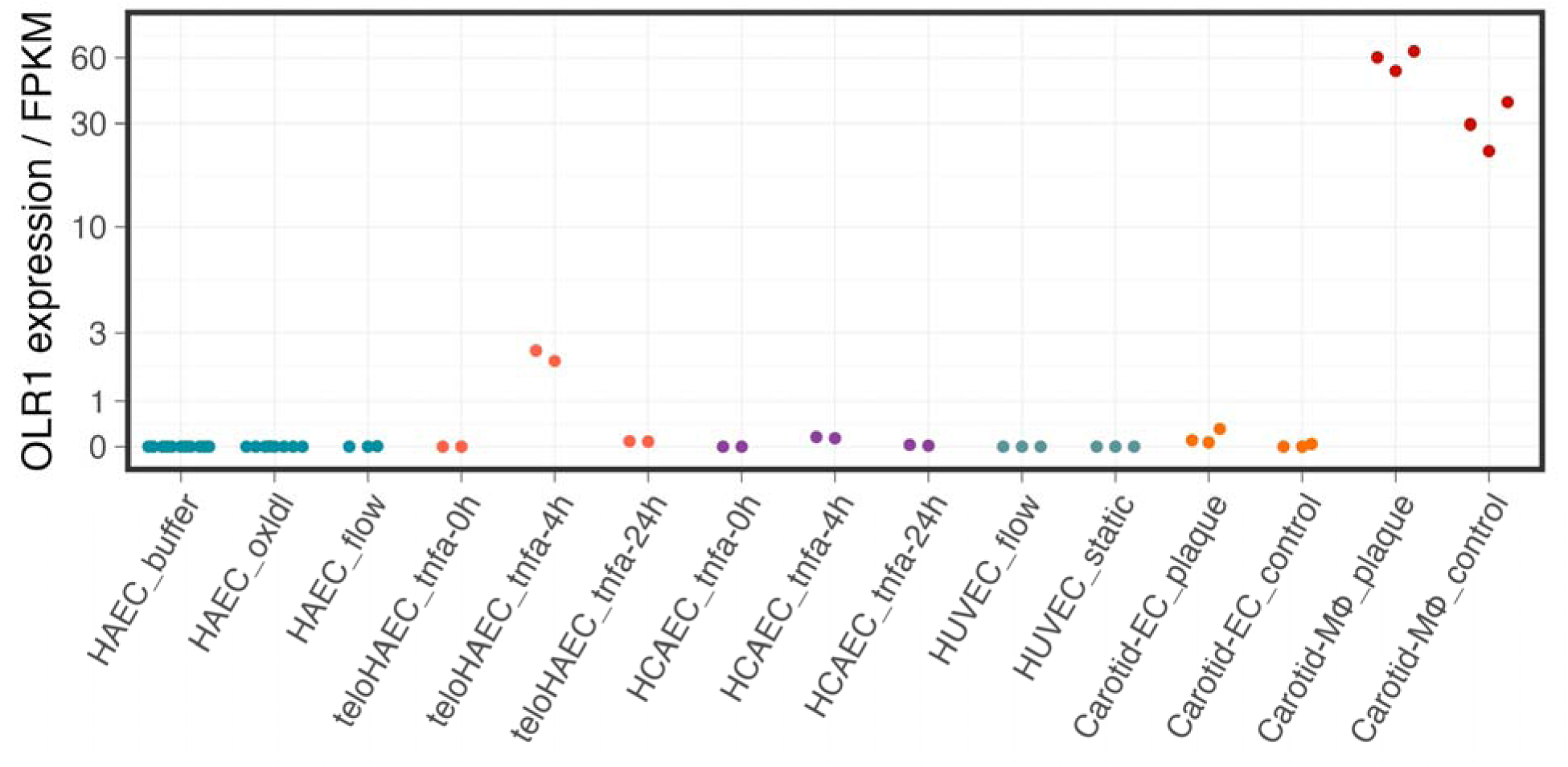
*OLR1* is lowly expressed in human endothelial cells under physiological conditions. The mRNA level of *OLR1* was examined across human endothelial cells of various origins including disease-free primary (HAEC) and immortalized aortic endothelial cell (teloHAEC) ^74^, coronary artery endothelial cell (HCAEC) ^74^, and umbilical vein endothelial cell (HUVEC) ^75^. Pseudo-bulk expression profiles of *in vivo* endothelial cells and macrophages from advanced carotid atherosclerotic plaque or adjacent disease-free tissue ^61^ are shown to the right for comparison. All HAEC datasets were generated in this study, including unpublished data on cells grown under laminar shear stress (HAEC_flow).

**Figure S3.**
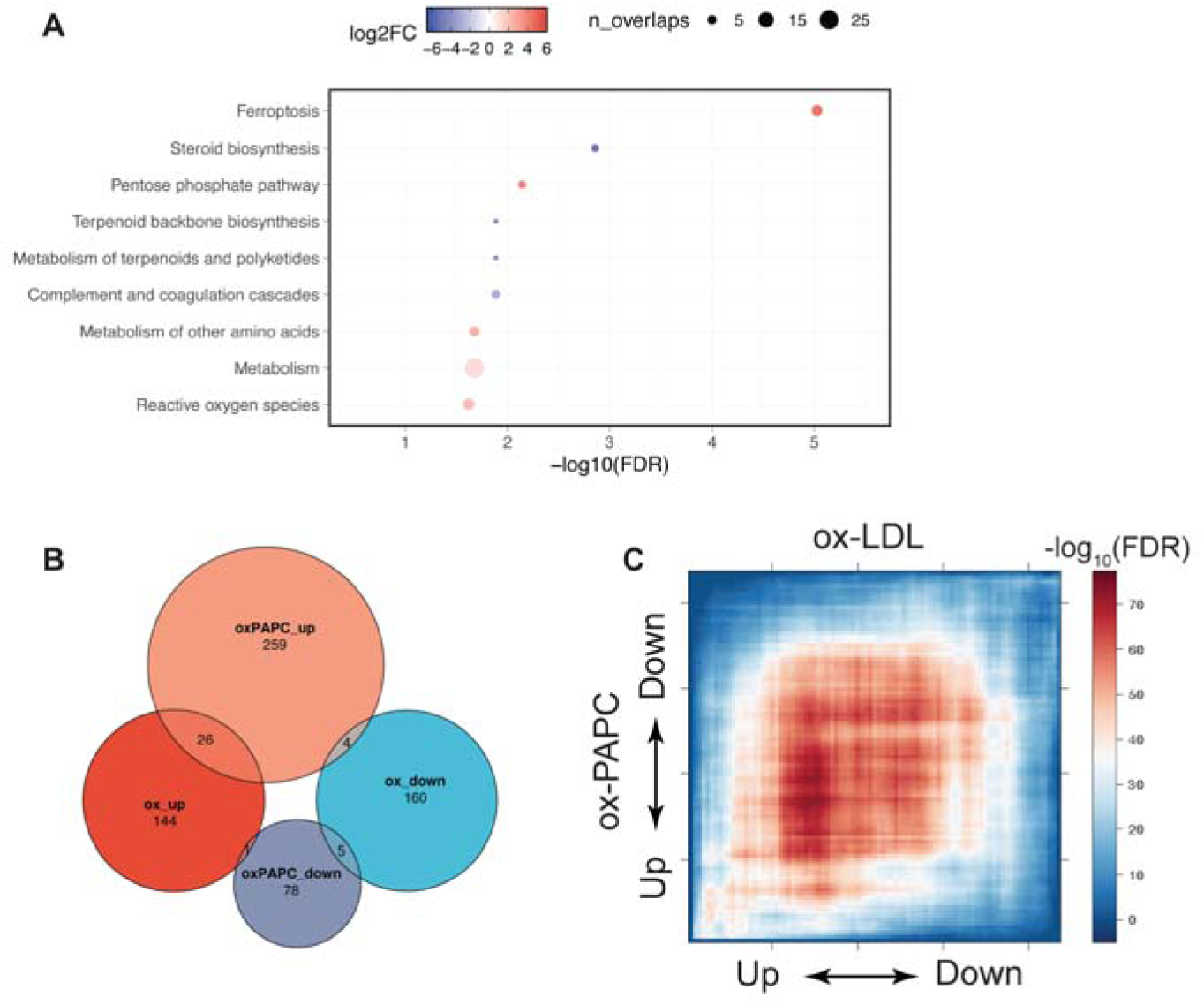
Limited overlaps between the transcriptomic response to ox-LDL and ox-PAPC in HAECs. **A**, KEGG pathways enriched in the differentially expressed genes upon ox-LDL exposure, ranked by statistical significance. Colour represents log2 fold change of pathway enrichment and size represents the number of overlapped genes in each pathway. **B**, Venn plot showing the overlaps between genes regulated by ox-LDL and ox-PAPC in HAECs. **C**, Rank-rank hypergeometric overlap tests showing the lack of overlaps in genes significantly regulated by ox-LDL (x axis) and ox-PAPC (y axis). The colour denotes the significance of each hypergeometric test (-log_10_FDR), measuring the extent of overlap between the two gene set on a sliding basis. Note the lack of significance at the end of x and y axis, indicating limited overlaps between differentially-expressed genes in the two datasets.

**Figure S4.**
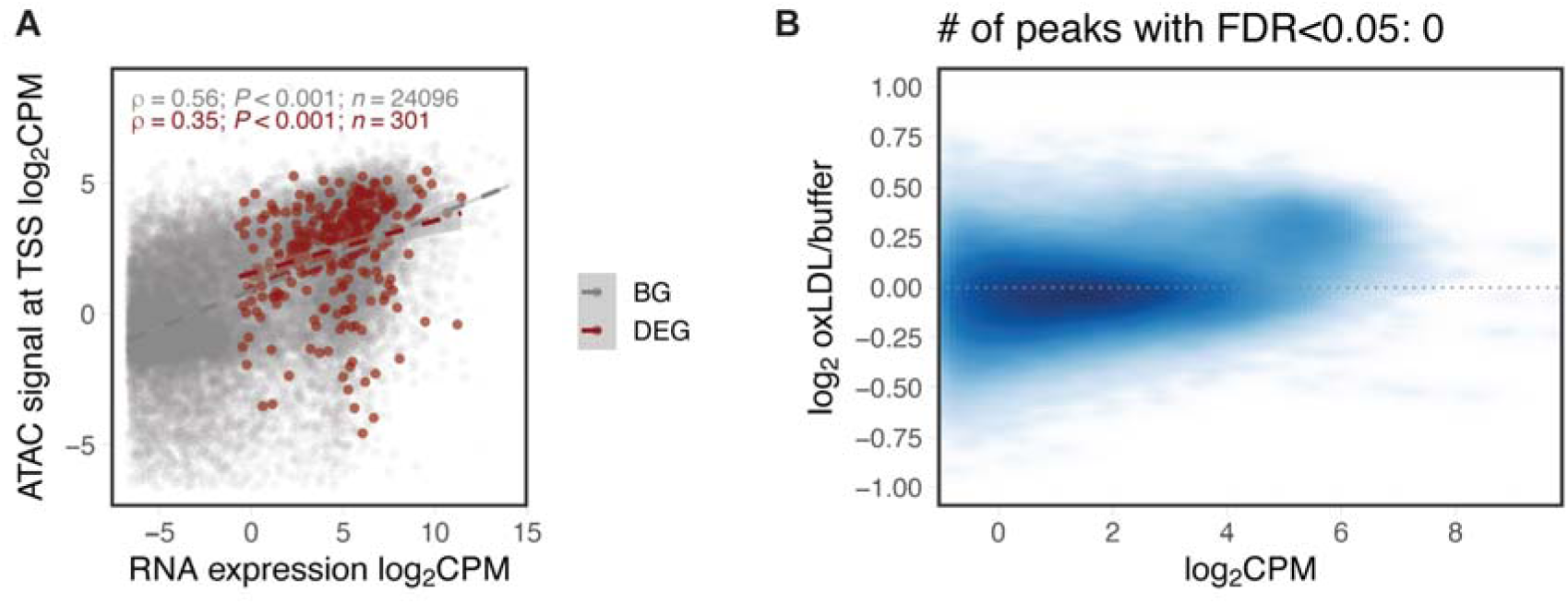
Ox-LDL does not induce global changes in chromatin accessibility in HAECs. **A**, Correlation between gene expression and chromatin accessibility at the 2-kb window centred at the transcription start site (TSS) in differentially expressed genes by ox-LDL (red) and all other genes (grey). **B**, Fold change in chromatin accessibility plotted against the average accessibility across all open chromatin regions in HAECs. No open chromatin peak is identified as differentially accessible under the FDR threshold of 0.05.

**Figure S5.**
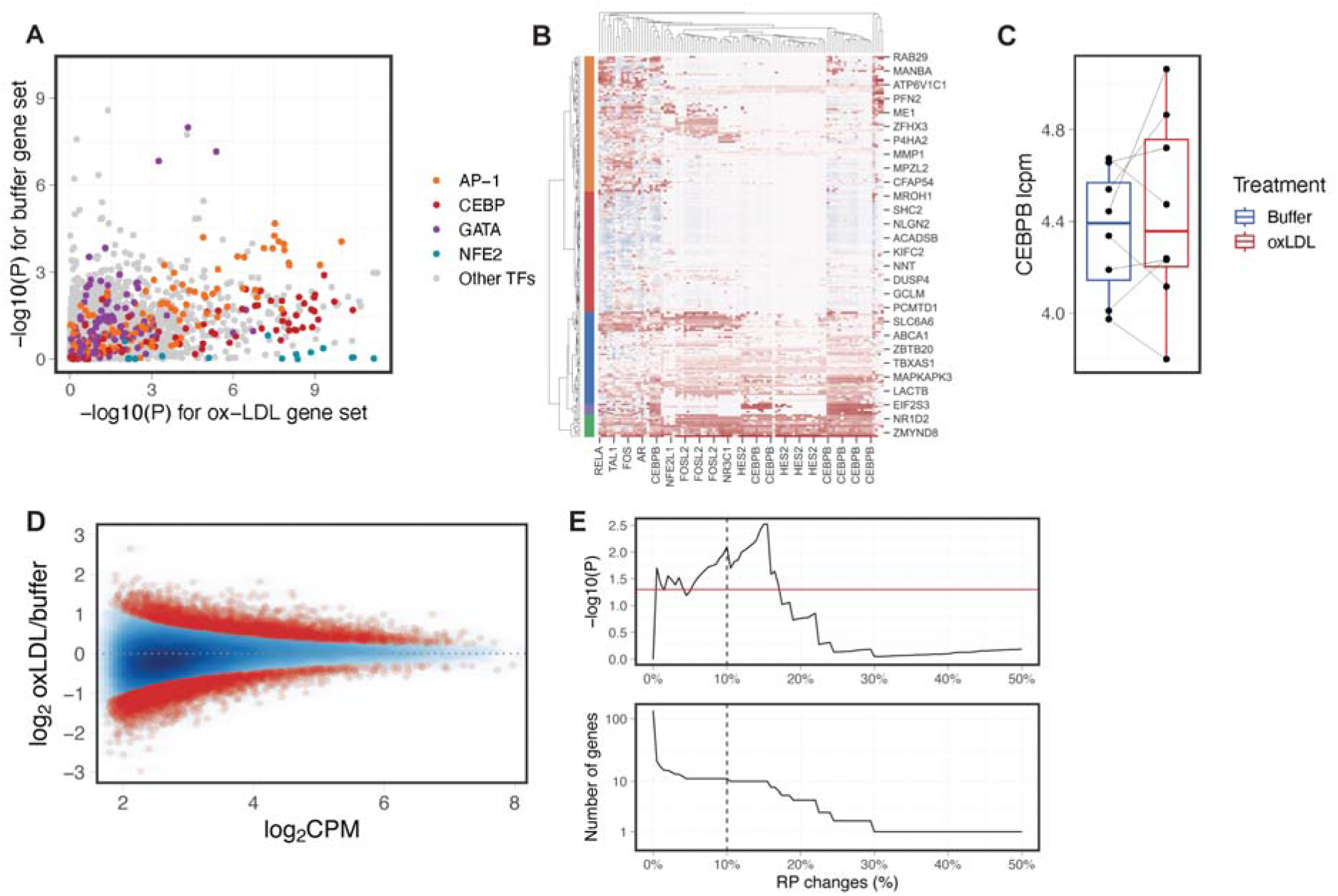
Ox-LDL-specific in-silico deletion analysis highlights CEBPB as a transcription regulator in HAECs induced by ox-LDL. A, ISD analysis was repeated using the enhancer landscape (H3K27ac ChIP-seq) instead of the open chromatin landscape (ATAC-seq), and p-values are shown similar to those in Figure 3D. **B**, Heatmap showing the predicted regulatory effect of each TF ISD (column) on the expression of individual genes (row) induced by ox-LDL treatment. The colour in in the heatmap represents predicted regulatory score (in an arbitrary unit), with red denoting activating effects. **C**, Normalised log-count-per-million (lcpm) gene expression of *CEBPB* as quantified by bulk RNA-seq, coloured by treatment group. HAECs were treated with 50ug/mL ox-LDL for 48 hours for RNA-seq experiments (n=4 biological replicates with 2 technical replicates per donor). **D**, Fold change in CEBPB binding plotted against the average ChIP-seq signals across all CEBPB binding sites in primary human macrophages. A subset of 8,718 differential binding sites under the p-value threshold of 0.05 (coloured in red) were selected for further ISD analysis. **E**, Enrichment test for genes disrupted by the ox-LDL-specific ISD in ox-LDL up-regulated genes. Disrupted genes are selected based on a sliding threshold of changes in regulatory potential, and the threshold used in Figure 3E is marked by the dashed line.

**Figure S6.**
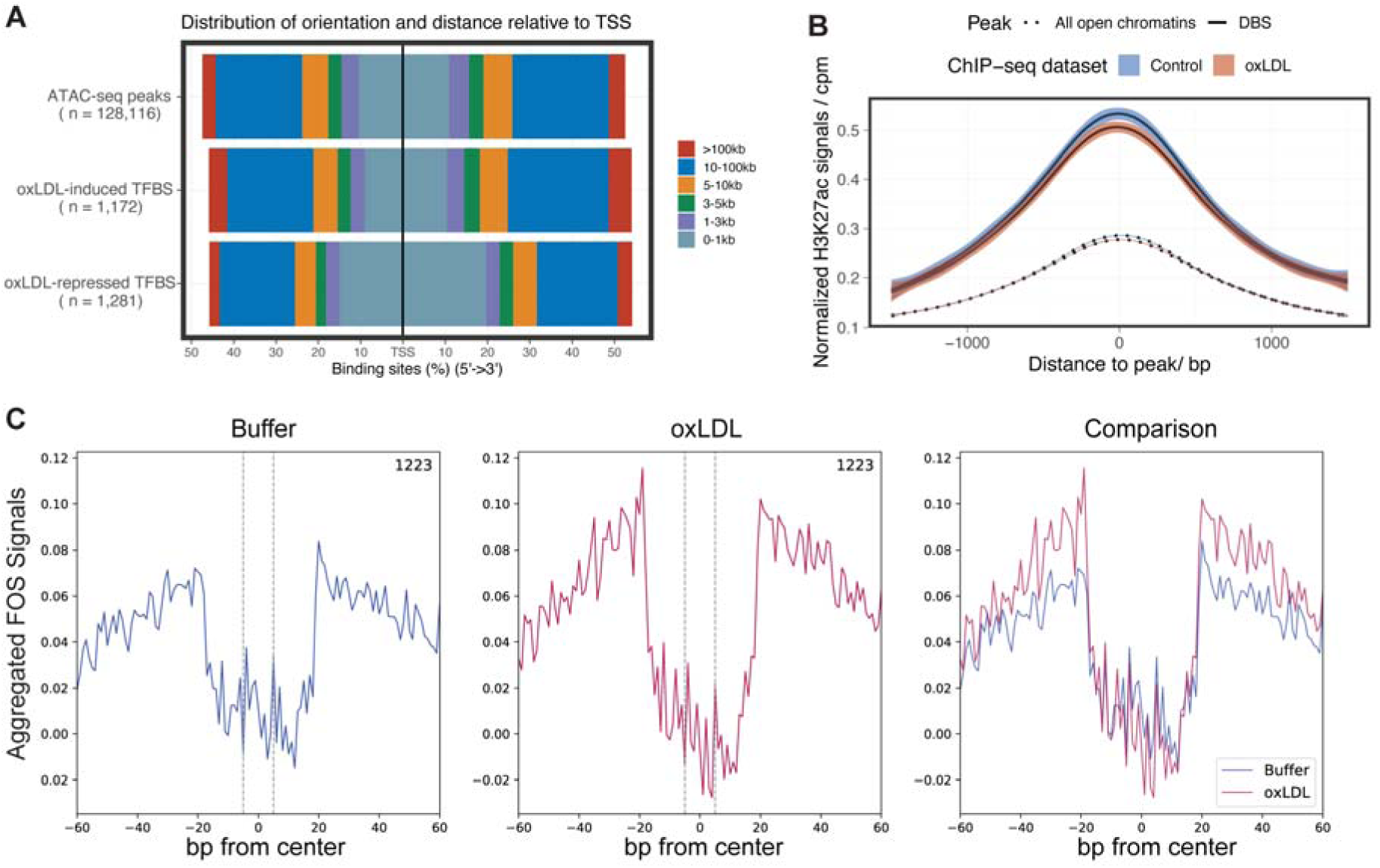
Motif footprinting analysis reveals redistribution of transcription factor binding in ox-LDL-exposed endothelial cells. **A**, Genomic annotation of the identified TF binding sites induced or repressed by ox-LDL. **B**, Aggregated H3K27ac signals at peaks containing dynamic TF binding sites (DBS) or background open chromatin regions. Signals from unexposed (blue) and ox-LDL-exposed (red) HAECs are plotted separately. Shaded regions indicate 95% confidence intervals. **C**, Aggregated foot print signals for FOS at n=1,223 potential binding sites with ox-LDL-induced activity.

**Figure S7.**
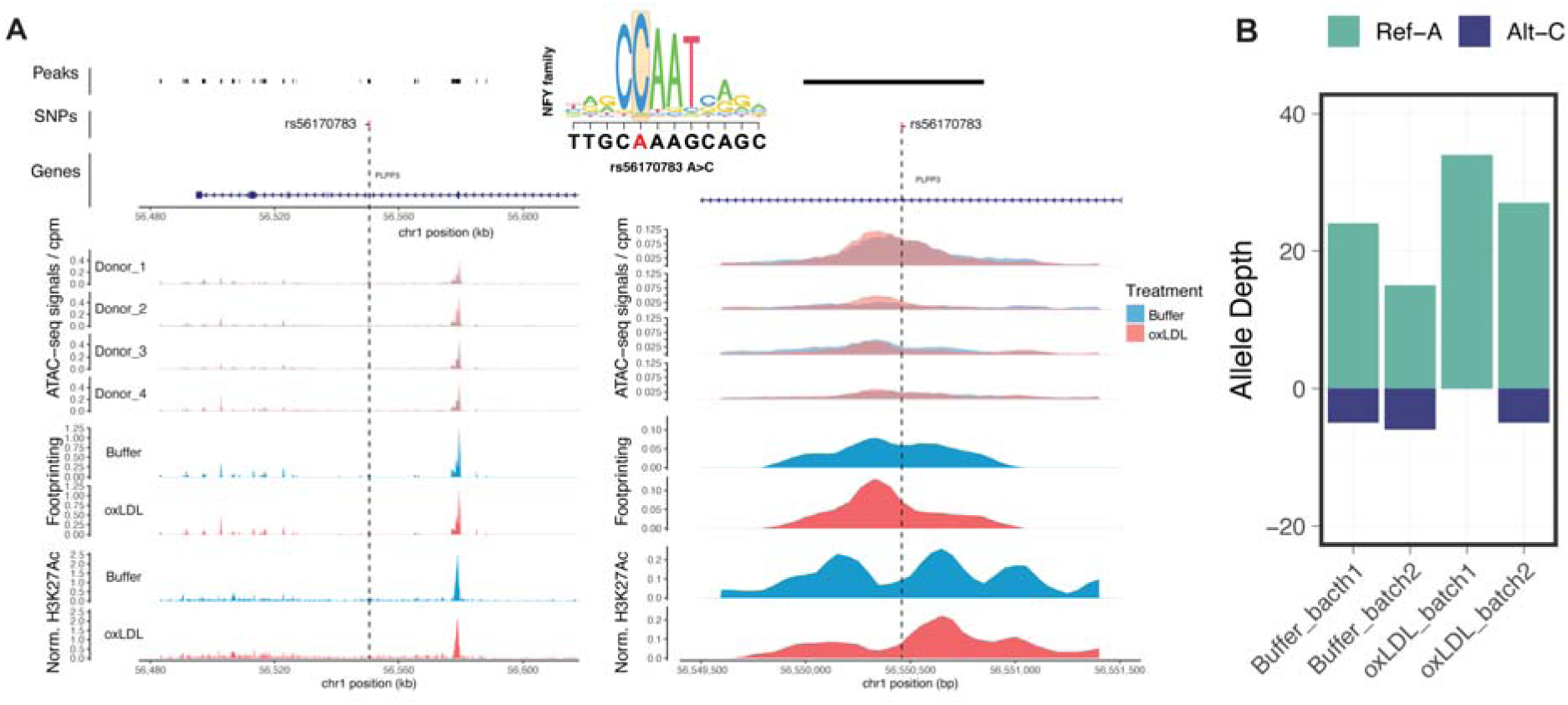
The regulatory landscape at the prioritised *PLPP3* locus. **A**, The candidate causal variant rs56170783 is marked in red. Tracks (from top to bottom) showing: ATAC-seq peaks (peaks), CAD-associated variants (SNPs), genomic annotations (Genes), ATAC-seq signals aggregated by biological donors with signals from paired samples overlaid together, Motif footprinting score aggregated by treatment, and enhancer signals from H3K27ac ChIP-seq. Higher resolution coverage plots centred on the risk variant are plotted in the right panel. **B**, Allelic sequencing coverage at rs56170783 in the ATAC-seq libraries of four heterozygous samples.

**Figure S8.**
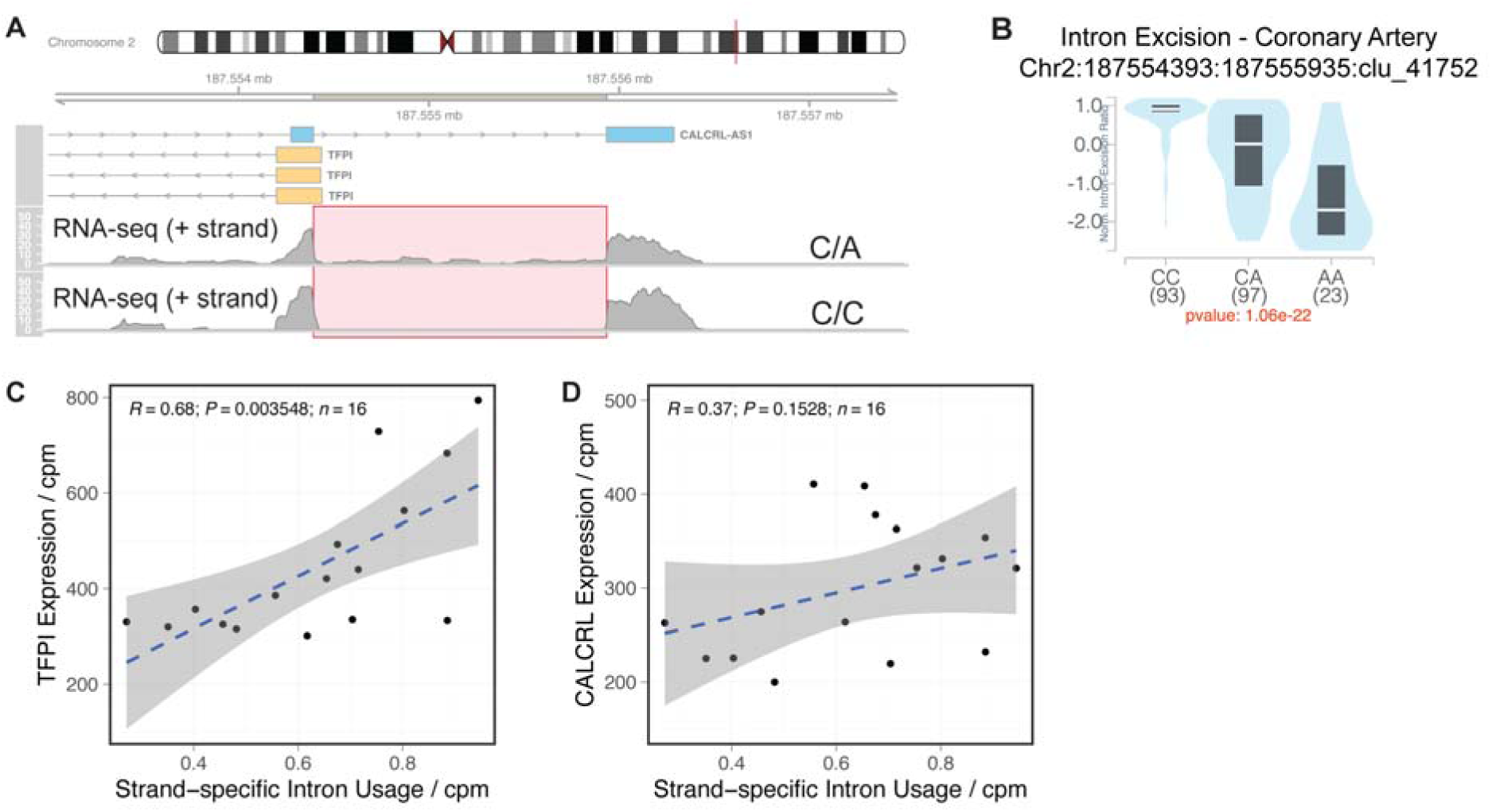
Alternative splicing of CALCRL-AS1 is associated with gene expression change in TFPI but not CALCRL. **A**, Overview of the alternative splicing event of the last intron of *CALCRL-AS1*. Coverage plots of RNA-seq reads on the positive strand are plotted in the bottom panel. The sample heterozygous for rs62172376 shows retention of the last intron, as highlighted in red, which is antisense to the promoter region of *TFPI*. **B**, Normalised excision ratio for the intron chr2-187554393-187555935 of *CALCRL-AS1* as shown in **A** in human coronary artery based on the GTEx data. Samples are grouped by the genotype at the SNP rs62172376. **C** and **D**, Pearson correlation tests between the strand-specific usage of the intron chr2-187554393-187555935 and gene expression of *TFPI* (**C**) and *CALCRL* (**D**). Gene and intron expression normalised to sequencing depth (count-per-million-reads). Shaded regions indicate 95% confidence intervals estimated from n=16 HAEC samples from 4 biological donors sequenced in this study.

